# Automated segmentation of insect anatomy from micro-CT images using deep learning

**DOI:** 10.1101/2021.05.29.446283

**Authors:** Evropi Toulkeridou, Carlos Enrique Gutierrez, Daniel Baum, Kenji Doya, Evan P. Economo

## Abstract

Three-dimensional (3D) imaging, such as micro-computed tomography (micro-CT), is increasingly being used by organismal biologists for precise and comprehensive anatomical characterization. However, the segmentation of anatomical structures remains a bottleneck in research, often requiring tedious manual work. Here, we propose a pipeline for the fully-automated segmentation of anatomical structures in micro-CT images utilizing state-of-the-art deep learning methods, selecting the ant brain as a test case. We implemented the U-Net architecture for 2D image segmentation for our convolutional neural network (CNN), combined with pixel-island detection. For training and validation of the network, we assembled a dataset of semi-manually segmented brain images of 94 ant species. The trained network predicted the brain area in ant images fast and accurately; its performance tested on validation sets showed good agreement between the prediction and the target, scoring 80% Intersection over Union (IoU) and 90% Dice Coefficient (F1) accuracy. While manual segmentation usually takes many hours for each brain, the trained network takes only a few minutes. Furthermore, our network is generalizable for segmenting the whole neural system in full-body scans, and works in tests on distantly related and morphologically divergent insects (e.g., fruit flies). The latter suggest that methods like the one presented here generally apply across diverse taxa. Our method makes the construction of segmented maps and the morphological quantification of different species more efficient and scalable to large datasets, a step toward a big data approach to organismal anatomy.

## Introduction

Three-dimensional (3D) imaging of animals by x-ray micro-computed tomography (micro-CT) has become popular in morphological biology as a non-destructive method to acquire high-precision data on organismal anatomy [1, 2, 3, 4, 5]. The high-resolution 3D data enables the users to visualize and quantify internal and external structures, forming the basis for a wide range of biological applications.

A key challenge for the use of micro-CT lies in the analysis of huge amounts of acquired data. In particular, while 3D images are reconstructed shortly after scanning, segmentation of the images into specific body parts is often a necessary step for quantification and visualization of particular structures. The most common segmentation method to date is by manual processing, which is extremely time-consuming and compromises reproducibility [6]. This limits the number of samples that can be included in a given study, and thus the scientific applications of 3D scanning. For example, developmental biologists may want to analyze large numbers of experimental treatments and replicates. Or, in comparative biology, we may seek to analyze the evolution of a body part across hundreds or thousands of species. The recent emergence of large databases and coordinated projects to scan many species in specific taxonomic groups offers rich opportunities for new research directions if limitations on segmentation can be overcome.

In the medical literature, image segmentation methods have recently become more powerful and efficient due to significant developments in machine learning algorithms. To date, the main focus of automated segmentation methods has been on cells and human organs (e.g., human CT or MRI image segmentation for cancer detection [7, 8]). However, there is broad potential for automated segmentation to accelerate biological research on organisms across the tree of life [9, 10, 11].

New software for biomedical image analysis has steadily progressed during recent years, with the capability for analysis and segmentation of 2D or 3D biological images and the capability to build one’s own data processing pipelines [12]. However, despite the unconstrained accessibility to free general-purpose software tools, the development of specific segmentation algorithms is essential to achieve high accuracy, objectivity, and reproducibility. Recently, deep learning and convolutional neural networks (CNNs) have been successfully applied in numerous image classification and semantic segmentation problems [13, 14]. CNNs have recently become widely used in image processing due to their high performance, the efficiency of GPUs, and the availablity of free software platforms and pre-trained networks [15].

Toolsets and pipelines that use classical statistical methods such as ANTs [16], Biomedisa [17], and Freesurfer [18] are accessible and accurate for the segmentation of high-resolution images. However, these are either not fully automated and still require an expert user and considerable amounts of time and effort [19], and/or require training examples within the same scan, and/or are not adaptable to diversity and complexity in the target set. On the other hand, accurate and general toolkits and application frameworks that use machine learning techniques such as SlideCam have been successfully used for medical image segmentation as well as computer-aided diagnosis and analysis of images spanning from human brain segmentation to cancer detection [20]. However, to date no toolkit has been designed to recognize homologous parts across a wide diversity of animal species, which would require an appropriate choice of network architecture, fine-tuning of hyperparameters, and the production and curation of substantial, high-quality datasets. When it comes to analyzing such images, segmentation remains a most challenging task, and often manual or semi-automated-segmentation is still the only way.

U-Net is a CNN architecture that has shown high accuracy and robustness for biomedical image segmentation [21]. It uses relatively small amounts of training images to achieve precision even for segmentation of areas with unclear borders. The simple architecture of U-Net makes it easy to develop and very fast to train. Once a U-Net is trained, the acceleration of the segmentation is extreme: for example, the segmentation time for one ant brain, which may be up to a whole day’s work if performed manually, is reduced to merely 1-2 minutes by automatic segmentation.

In this paper we present an automated pipeline for segmentation of different inner parts of insects in volumetric data, using micro-CT scans, and specifically ant brains across a diverse set of different ant species, as a test case. A basic question for such studies is how general algorithms can be applied across the tree of life. Can an algorithm trained to recognize a part in one type of organism be used on more distant relatives, or do they break down once applied outside the group for which they were developed? Ants are a well defined clade following a similar overall body plan, but reflect > 100 million years of diversification and a large range in ecological, sensory, and behavioral modes [22, 23]. We expect ant brains to have an intermediate level of diversity and thus be a reasonable test case: they will change in size and shape across species, while the general organization and tissue composition should be conserved [24]. As a secondary experiment, we assess whether the ant brain segmenting algorithm we developed can be applied with minimal modification to recognize brains in distantly related insects.

### Overview of the segmentation pipeline

Our micro-CT image segmentation pipeline is composed of multiple modules, as illustrated in Fig 1.

**Fig 1.**
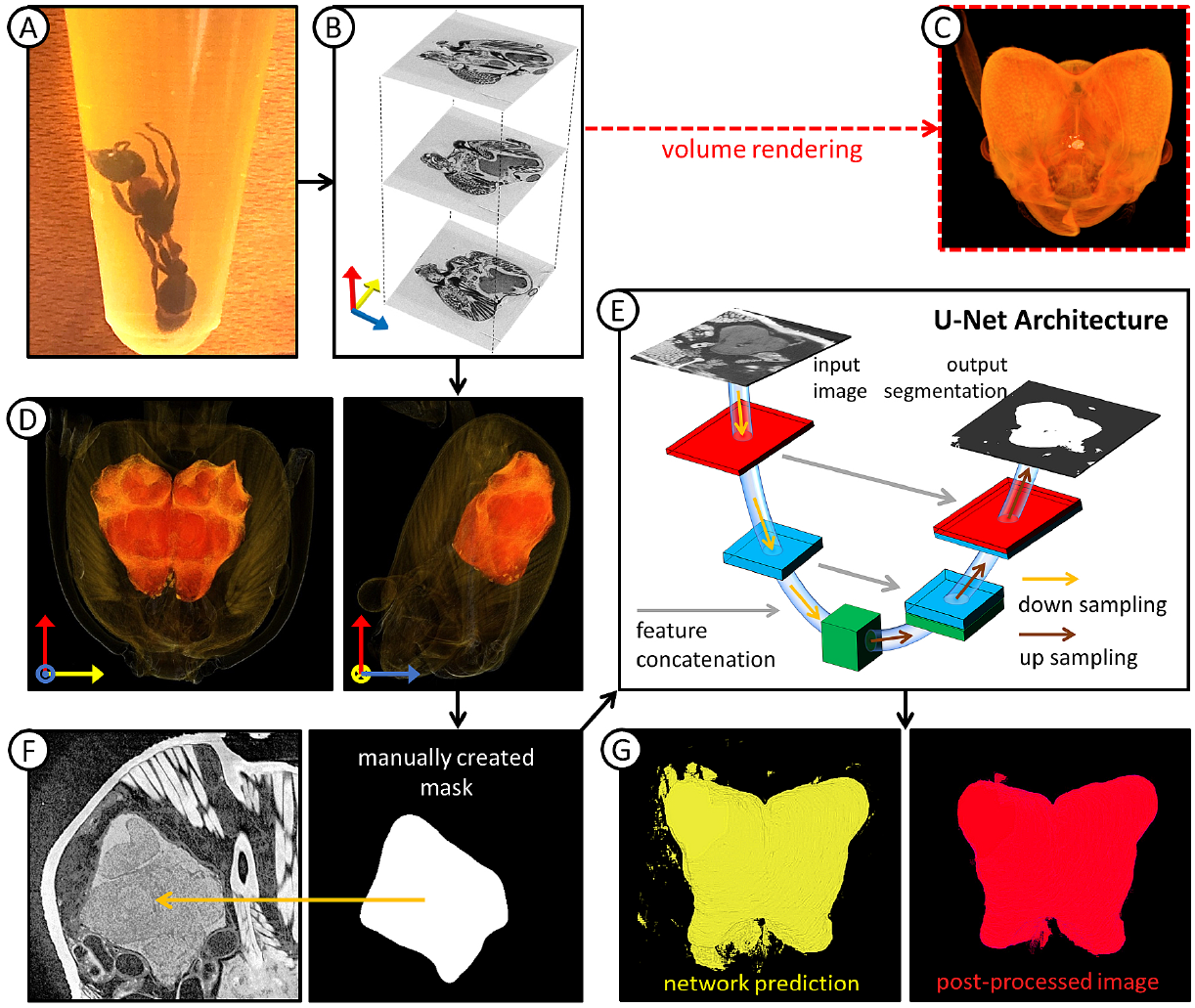
Segmentation pipeline overview. (A) Specimens are placed in iodine for staining for two weeks and then placed in small vials containing 99% ethanol to prevent them from moving during scanning. (B) The scanner acquires images along all three axes, and, using a user-defined reference image, automatically reconstructs the whole volume of the scanned specimen. (C) Volume rendering for future morphological studies is performed using Amira software. (D) Semi-automated segmentation of the brain volume of each scan (in orange) using the watershed method in Amira. (E) Schematic representation of the U-Net architecture used as the core of the pipeline for the development of a fully automated brain segmentation method. (F) The acquired brain images are used for training after pre-processing augmentation and manual creation of masks. (G) The network’s prediction (in yellow) is post-processed for smoothing out over-predicted areas (in red).

- **Sample preparation:** Before scanning, specimens were stained in iodine for an average of two weeks to enhance tissue contrast in the raw images.
- **Image acquisition and reconstruction:** An X-ray micro-CT image dataset was acquired from 76 species of ants. The acquired images were reconstructed along all three perpendicular directions that comprise a Cartesian system forming a detailed cross-section dataset.
- **Volume rendering:** The reconstructed raw images were used for creating a 3D model for volume rendering, to be used for visual inspection and future morphological studies.
- **Semi-automated segmentation:** Raw images of heads were segmented semi-automatically using the seed-based watershed tool of the Amira software. Labels were assigned to areas of interest, starting with the brain. The databases of both raw and labeled images were pre-processed to enhance their homogeneity and used as training and validation data.
- **CNN development:** An implementation of the U-Net architecture was built for automated segmentation.
- **Training:** 60% of the acquired segmented brain images (46 species) were used for network training; the remaining 40% (30 species) was reserved for testing.
- **Pixel island detection and post-processing:** After segmentation by U-Net, pixel island detection was used to identify the largest continuous areas to remove isolated segments.

## Materials and methods

### Image acquisition

In total, we collected one head scan per species from 76 different ant species using a ZEISS Xradia 510 Versa 3D X-ray micro-CT microscope, and ZEISS Scout and Scan Control System software (version 10.7.2936). The scanner settings were determined by the specimen size (e.g., voltage: 30 keV and exposure time: 3-10 s) resulting in 5-to 20-hour scans (12 hours on average). With a view to expanding our dataset in order to eventually enhance robustness and suppress overfitting during network training (see below), we used 2D cross-sections of planes along all three directions of our 3D brain scans. To highlight the morphological diversity of the scanned specimens, we also performed 3D reconstruction of the resulting scans with XMReconstructor (version 10.7.2936). The output images comprised 1000×1000×1000 px, on average, with resolution down to 1 *µ*m. Exemplary raw images of full-body scans from different ant species are shown in Fig 2.

**Fig 2.**
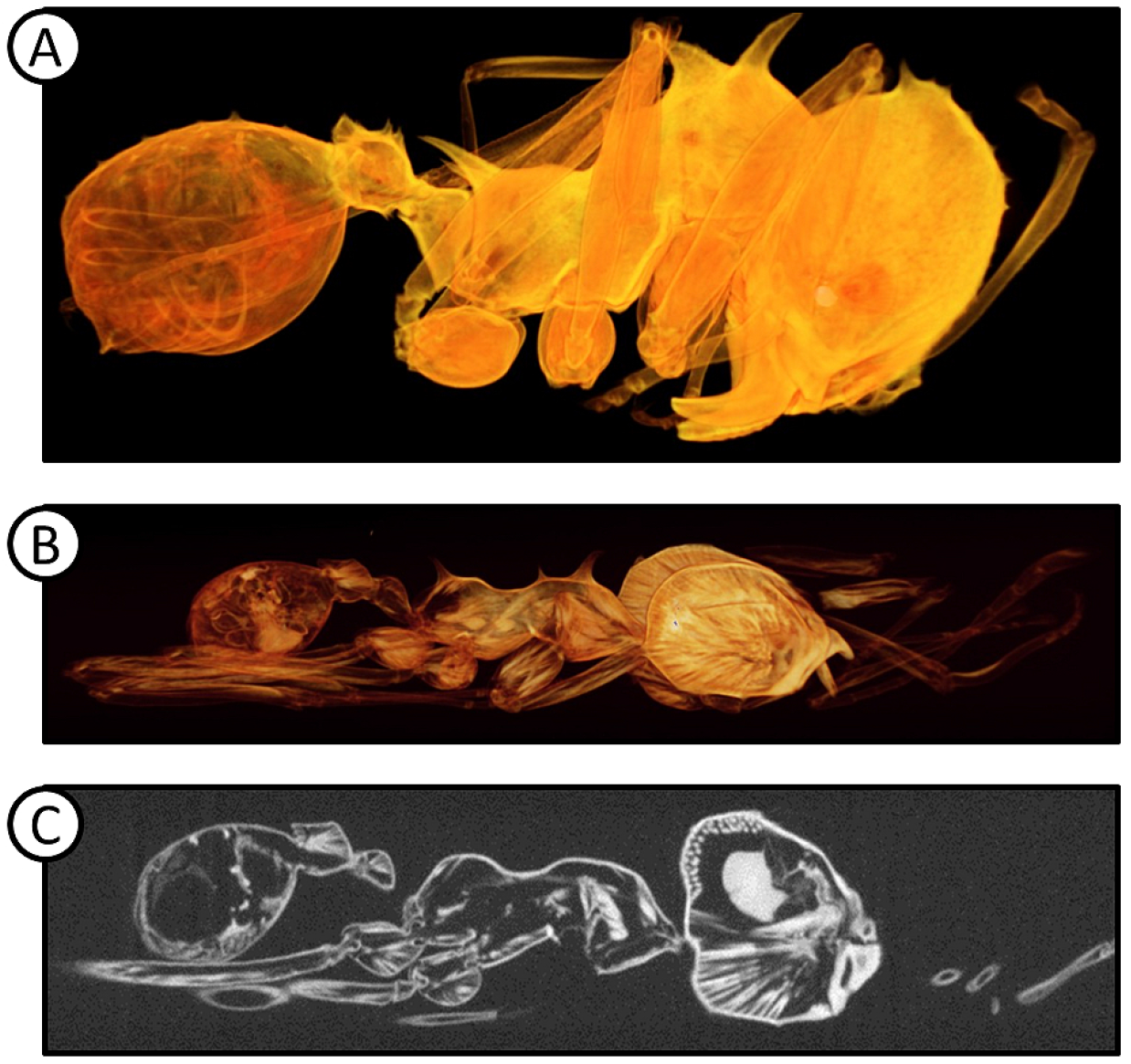
Exemplary raw images of full-body scans from different ant species. 3D reconstructed micro-CT image of (A) *Acromyrmex versicolor* and (B) *Atta texana* worker specimens, using volume rendering in Amira. (C) 2D micro-CT full body image of *Atta texana* specimen. The brain area is the densest, most uniform area in the whole body, which makes it easy to recognize in most high-quality scans.

### Image processing

#### Generation of data for training and validation

We processed the data with the Amira software (version 6.0), and semi-automatically segmented the brain areas (on average, 300×400×600 px per brain) using the seed-based watershed method [25] in the volumetric data, as shown in Fig 3. Each semi-automatically segmented 3D brain was dissected into 2D slices on planes along all three directions. To eliminate the empty space of the image and zoom in on the region of interest, the image was cropped and rescaled in to 500×500 px, to show only the brain area and its adjacent muscles and fibers. The texture of the brain is unique within the whole image of the head, which facilitates its identification. However, its borders are much harder to classify, as numerous nerves branch out from the brain connecting it with the rest of the ant’s body; these nerves had to be removed manually, as Amira’s watershed tool typically mistakes them for brain areas. This makes semi-automated segmentation challenging and considerably time-consuming. Eventually, this process resulted in an average of 1000 2D brain images per specimen at an estimated average time cost of 5 hours per specimen.

**Fig 3.**
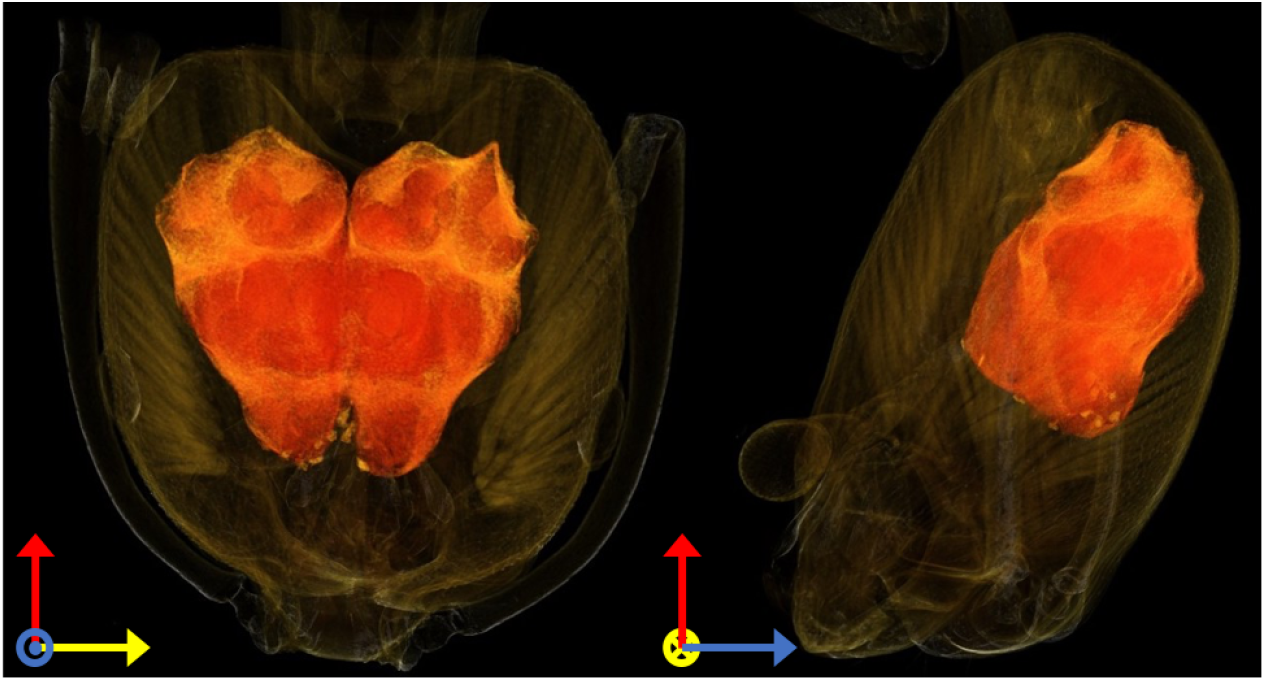
Example of semi-automated brain image segmentation. The brain area (in orange) of an *Atta texana* ant specimen was segmented using the watershed method in Amira; the image was manually post-processed by smoothing and cropping over-segmented areas.

### Pre-processing and augmentation of data

As a pre-processing method, we chose histogram equalization [26] using the imaug library [27] in Python. Since the images were collected from different samples and scans, and their contrast was not optimized during scanning, pixel density equalization improved the network’s performance remarkably. After histogram equalization, the contrast was improved, accentuating the texture of the brain and, thus, making it easier to identify. An exemplary result of data augmentation is shown in Fig 4.

**Fig 4.**
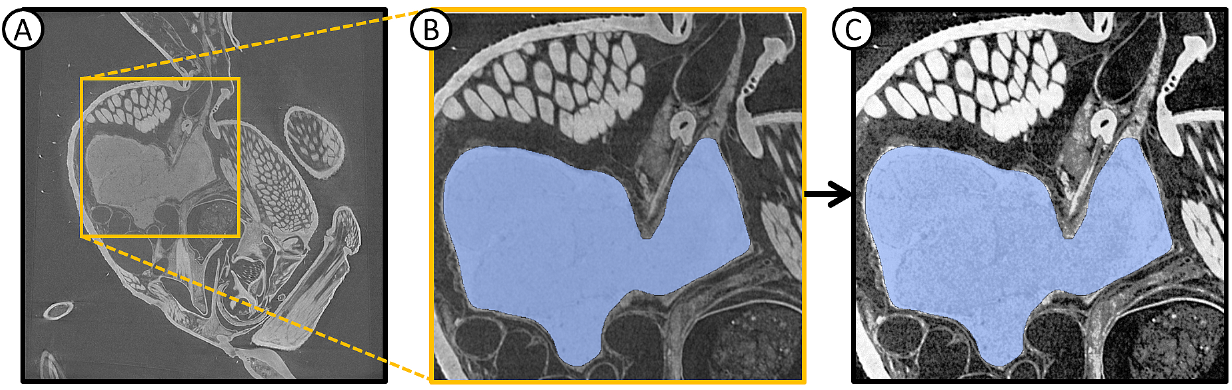
Data augmentation. (A) Initial 2D image of a full head scan of an *Atta texana* ant specimen. Pre-processing is performed in two steps: (B) The image is cropped around the brain area, keeping some of the muscles, nerves, and fibers that are close (or even attached) to the brain. The manual segmentation of the brain is indicated in blue. (C) Histogram equalization is used for additional augmentation, which enhances the contrast and projects the inner parts of the brain more clearly.

#### U-Net structure

We chose the U-Net architecture because it has been the most successful CNN for CT image segmentation to date. The U-Net is not a conventional CNN architecture, in the sense that it extends the contracting path of a typical CNN by a symmetrical expansive path [21]. For optimal efficiency, our code uses the open source GPU-TensorFlow library [28] and the TensorFlow U-Net implementation, as described in Akeret et al. [29], utilizing Jupyter notebook and Python. Our network consists of a five-fold repetition of two 3×3 convolutions followed by a rectifier linear unit (ReLU) and a 2×2 max pooling. Starting with 64 features, each layer doubles their number resulting in 1024 features before starting the expansive path which consists of two 3×3 convolutions followed by ReLU and 2×2 up-convolutions. The number of features is halved with each up-convolution but the result is concatenated with the features from the matching contraction layer. Finally, 1×1 convolution is applied to map each feature to our number of classes, i.e., two. In total, the architecture consists of 23 convolutional layers. The batch size used was 1∼4, the stride was 1, and zero padding was used for the max pool.

#### Training

To assess the effect of various parameters on the performance and the processing time, different batch sizes, numbers of initial features, epochs, and iterations were tested. We randomly selected 60% of data (46 species) for training and used the rest (30 species) for testing. The optimal parameter values were chosen on grounds of low computational cost and high classification accuracy for training data. We trained the network by optimizing the binary cross-entropy function with L2 regularization using stochastic gradient descent with the momentum of 0.8. The initial weights were selected by using a Gaussian distribution, in agreement with Ronneberger et al. [21]. Batch normalization was added in the first 3 layers to avoid overfitting as well as to accelerate training [30]. Finally, we added dropout in the first 3 layers equal to 0.5 also to avoid overfitting. We trained our network for 10 epochs, with mini-batch 32 on a 520×520 pixel image, costing 120 hours in our workstation using a GeForce GTX TITAN Xp and a GeForce GTX 1080 graphics cards.

#### Post-processing

We post-processed our network’s prediction by using pixel island identification and isolation [31]. After predicting the brain area along all three planes, the biggest pixel island was chosen as the brain area. This process boosted by almost 10% on average our prediction success rate of both the Jaccard Index (IoU) and Dice Coefficient (F1 score) [32].

## Results

### Segmentation of ant brains

First, we applied our method to our primary taxonomy group of choice, i.e., ants, and trained our network to segment the brain areas in micro-CT scans from different ant species. Our processed data of 38,000 520×520 pixel images from 46 species were used for training and validation (randomly split into 80% for training and 20% for validation) and the remaining 20,000 520×520 pixel images from 30 species were used for testing. As shown in Table 1, both IoU and F1 scores were steadily increased as we added more 2D images from planes along the same x-y directions of different species, and even more so after we included reconstructed 2D images from planes along all three directions of our 3D brain scans. To estimate the generalized performance of our network, we calculated the true positive rate (TPR) values and false positive rate (FPR) values of our images by changing the discrimination threshold of our network [33], shown in Fig 5. All values are close to 1 for both training and testing sets while the false positive rate (FPR) values remain less than 0.4 for most cases, indicating that our network predicted the brain region and its border accurately but without over-predicting. Results for test and training images are similar, suggesting good generalization capabilities for optimized hyperparameters of our network.

**Table 1.** Performance evaluation of our proposed pipeline. Both performance descriptors studied (IoU and F1 scores) increase steadily with increasing number of images and post-processing.

| Accuracy scores |  |  |
| --- | --- | --- |
| Number of images of training set | IoU | F1 |
| 3,500 - xy plane | 50% | 62% |
| 10,000 - xy plane | 63% | 71% |
| 38,000 - along all three directions - no post-processing | 72% | 80% |
| 38,000 - along all three directions - after post-processing | 80% | 90% |

**Fig 5.**
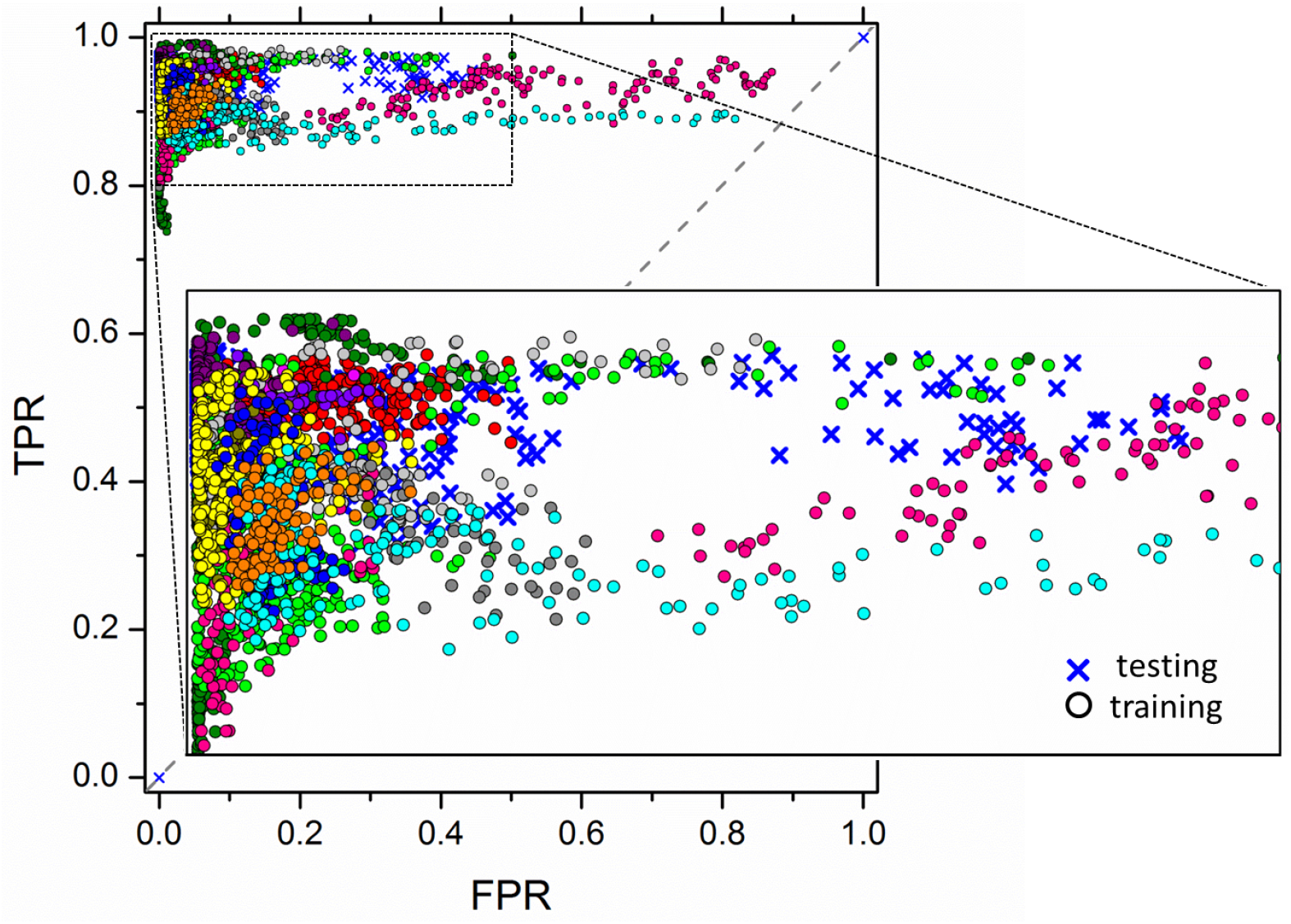
Network performance evaluation. High TPR and low FPR values for training (circles) and testing data (crosses) indicate the network’s high generalizability. Different colors correspond to brain images from different species.

Finally, a post-processing step also boosted the performance of our network, yielding even more satisfactory results. Example results of our network’s performance on validation and testing data are shown in Fig 6, demonstrating a predicted area in good agreement with the ground truth; our automated segmentation pipeline achieves an approximate maximum of 80% IoU and 90% F1 score. Prediction times were in the order of only a few minutes, significantly lower than for the semi-automated segmentation commonly used to this day.

**Fig 6.**
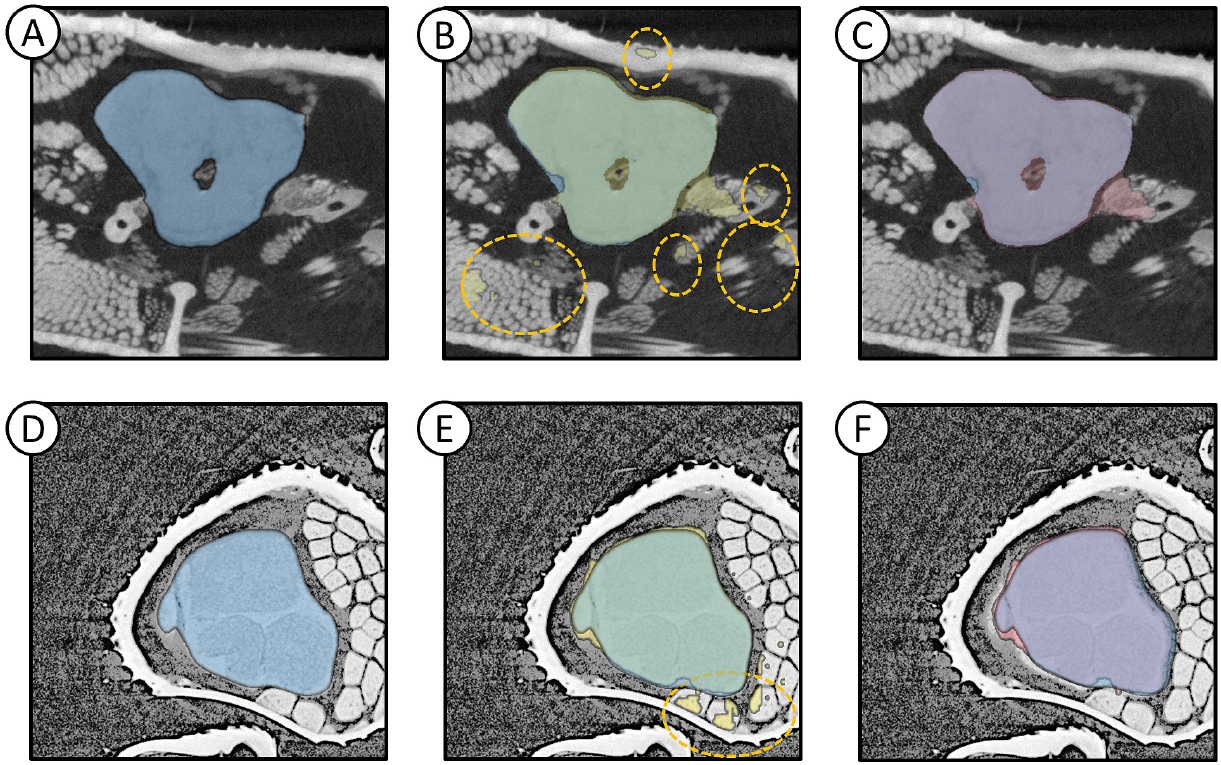
Pipeline performance calculated both for validation (top row) and testing (bottom row) sets. (A, D) Raw images of head of *Atta texana* ant and *Carebara atoma* specimens, cropped along the x-y axes. The manually segmented brain areas are indicated in blue. (B, E) Network predictions before post-processing (in yellow). Areas in yellow dotted circles are pixel islands not connected to the brain area that were overpredicted. (C, F) Predictions after post-processing (in red). The borders of the predicted areas show good agreement with the manual segmentation in both sets. Note that in overlapping manually and automatically segmented areas in B, C, E, and F, colors appear green or purple.

### 3D volume rendering

After segmenting the 2D slices, the 3D brain volume was readily computed by loading the stack of images in Amira or ITK-snap. Thus, using a 2D network allowed us to maintain high accuracy, performing 3D segmentation in a faster and easier to train way. An exemplary predicted brain area is shown in Fig 7; 3D volume was reconstructed from the 2D predicted images with Amira software. The switch from 2D to 3D is straightforward, giving the user of our pipeline the ability to adapt it to their own dataset circumventing the complications of using an actual 3D CNN.

**Fig 7.**
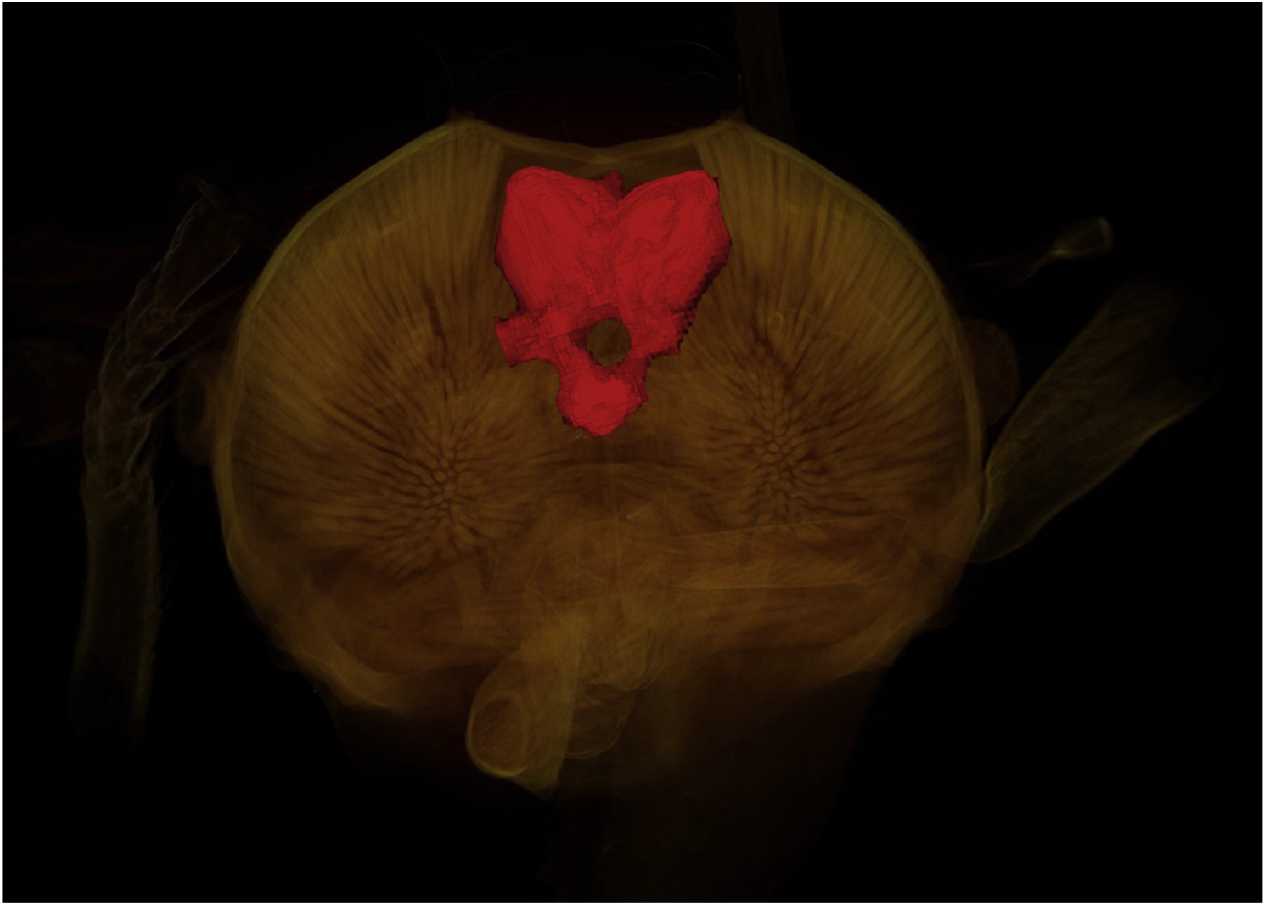
3D volume of ant brain reconstructed from 2D predicted images using the Amira software. 3D reconstructed brain prediction of *Atta texana* ant.

### Generalization to other neural systems and other insects

The U-Net step appears to be largely driven by textures, with the pixel island detection step used to isolate the brain. Even though our customized U-Net was designed with the segmentation of ant brains, it was also successfully applied for the segmentation of neural tissue in other parts of ants and works on distantly related species. Our network was able to predict the whole neural system in full-body scans of ants, as shown in Fig 8, being able to predict the same texture as the brain in different ganglia in the thorax (called mesosoma in ants).

**Fig 8.**
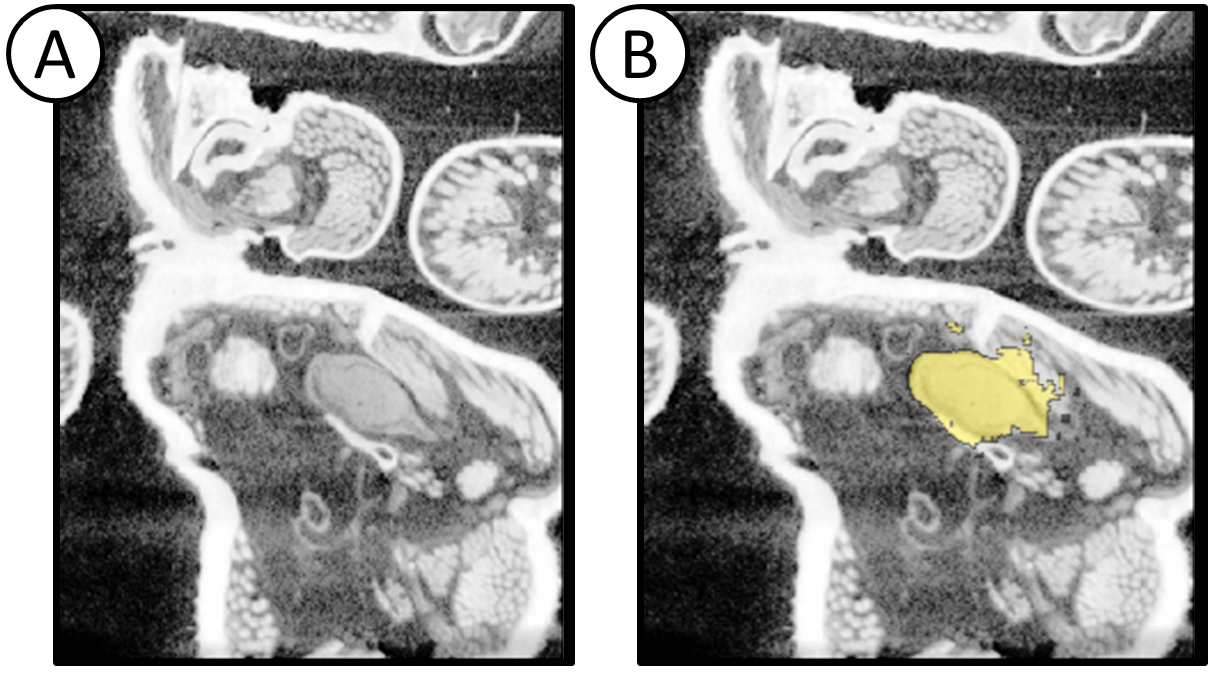
Prediction of ganglia in the thorax. As the tissue texture in the image is similar with that of the brain, the network accurately predicts other areas of nervous tissue in the organism. The pixel island detection step isolates the brain, but without this step neural tissue can be isolated.

Our network also gave good prediction for the brain area in scans of various different distantly related insect species. We used our pre-trained (on ant-brains) network to segment the brain areas of micro-CT scans of model organisms such as flies (*Drosophila*) and honey bees (*Apis mellifera*), as well as closely related insects such as praying mantises (*Leptomantella*) and termites. Since its prediction capability relies mainly on identifying the texture of the brain area, which does not differ significantly among different insect species, our pre-trained network was able to perform satisfactorily without further adaptation on the data. Exemplary results are shown in Fig 9 for (A-B) wasp and (C-D) praying mantis brain prediction, respectively: remarkably, our network was successful in segmenting the brains of different insects without any prediction accuracy losses (when compared to predictions for ants), indicating its flexibility and its lack of necessity for training on each specific distinct species.

**Fig 9.**
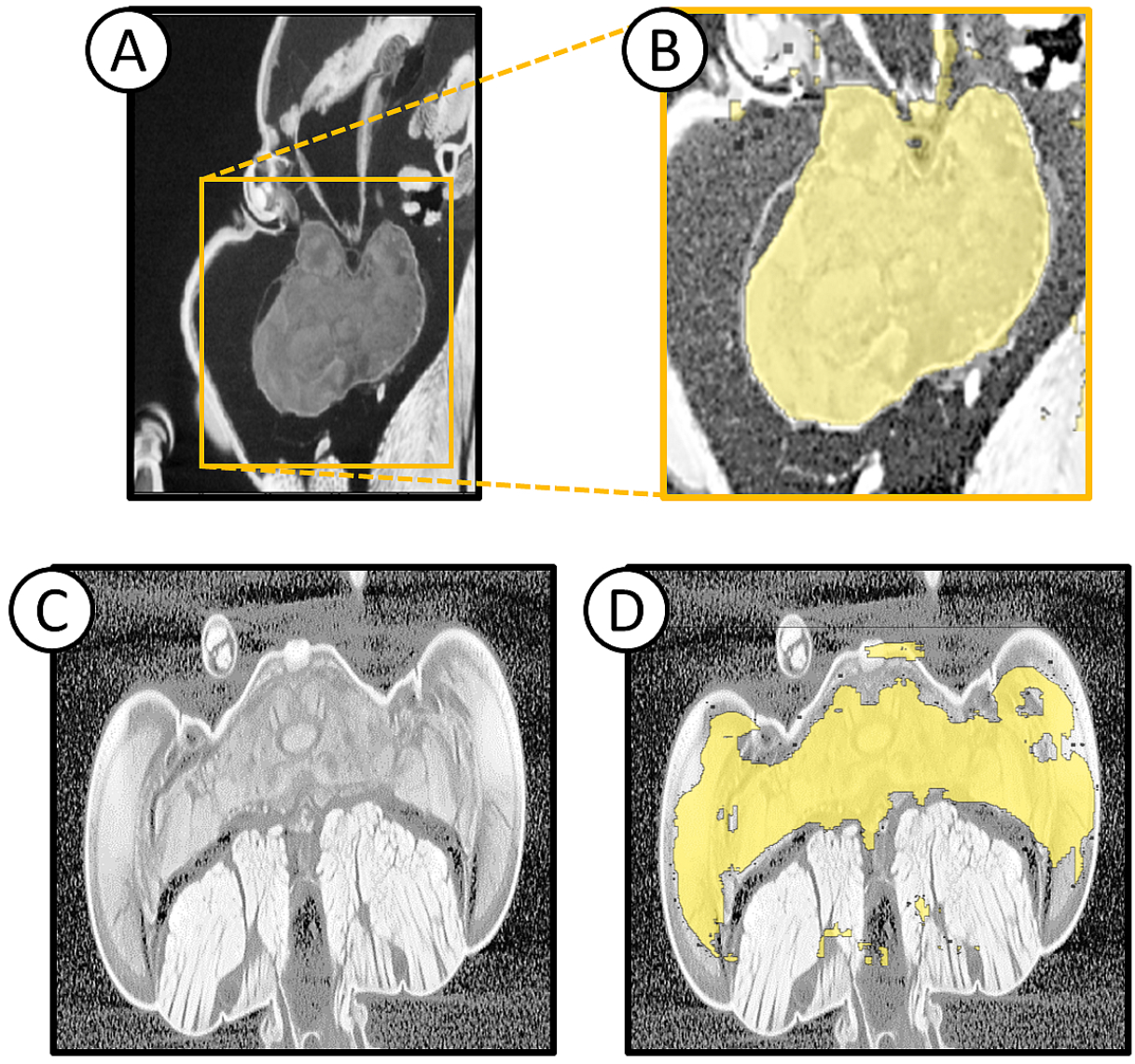
Application of pipeline for other insect species. The brain textures of various insect species can be very similar with those of ants, facilitating the prediction by the network even without pre-training on specific insect brain scans. (A) Raw image of wasp head and (B) its prediction without post-processing, indicating satisfactory identification of the borders of the brain area. (C) 2D image of praying mantis head and (D) the prediction of its brain area without post-processing. Even though the network over-predicts some small pixel islands, it excludes from its prediction areas of the muscles, fibers and cuticle.

## Discussion

To bring morphology fully into the big data era, we need automated methods to retrieve biological meaning from large volumes of images. The proposed automated pipeline is a step in that direction, presenting considerable advantages over other standard methodologies. First of all, automated segmentation is achievable within a few minutes for each specimen, producing faster and more accurate results than semi-automated or manual segmentation. A noteworthy additional advantage is that once algorithms have been trained, advanced expertise in morphology is not required, while manual and semi-automated segmentation usually require advanced knowledge [34]. In fact, during testing our network often outperformed even experienced users and compensated for their oversights or misjudgments, predicting correctly brain areas that were accidentally missed out during manual segmentation.

The two approaches in our method, U-Net and pixel-island detection, represent two complementary steps which suggest a path forward for automated segmentation of structures in complex organisms. U-Net was efficient at retrieving tissue with similar properties in the image, but in our implementation did not make use of shape and position. Thus, we found it retrieved all the structures of neural tissue across the body, even though it was trained on the brain alone. The brain was then isolated with the pixel-island detection, which isolated the largest structure in the head. In general, we expect a combination of tissue-level identification followed by other methods that make use of size and spatial organization to be a powerful combination that should generalize to a wide range of anatomical tissues and parts.

During testing with other insect species, we used both high and low resolution/quality images acquired from different laboratory and synchrotron-based micro-CT scanners. Our results showed that our segmentation pipeline can perform without losing its accuracy to predict the brain area across highly divergent arthropod species and across scanning methods. Finally, the prediction performance of low-resolution images indicates that there is a threshold in the image resolution below which our network is not performing well. Our network’s generalizability is high and it can be widely used not only for head but also for whole-body scans of ants and other insects. More importantly, it shows that a similar approach could be used to build a suite of trained networks that can segment anatomy across a wide variety of organisms.

Last, it should be noted that both automated classification and segmentation tasks typically require big datasets for training and validation, which can be a challenge for researchers to produce for any given application. Since no publicly available dataset of micro-CT images of ant brains existed for our case study, we created a new, extensive dataset across a wide variety of ant species. Since neural anatomy across insects share features that make them targets for segmentation, our dataset can act as a starting point for the development of an even bigger library of micro-CT images of insects, and work as a pre-training dataset for future CNNs [35].

## Conclusion

In this paper, we introduced a U-Net based CNN for the fully-automated segmentation of micro-CT images of insects. We also present an extensive dataset of manually segmented brain images that can be used to pre-train other networks of interest. Our trained network predicted the brain area in ant images fast and with high accuracy. Further, our network was able to generalize and predict the whole neural system in full-body scans, as well as to predict ganglion areas that were missed by manual segmentation. After training, the network’s performance was tested on training and validation data showing good agreement between prediction and mask scoring 90% F1 and 80% IoU. Our pipeline allows successful segmentation in only a few minutes instead of hours which are typically required for manual segmentation.

One of the most important features of the framework described here is that it can be applicable to other anatomical features. Preliminary results on other organs have shown that it can be easily tuned and trained to predict muscles as well as the cuticle of the insect bodies. Specific attention was paid so that the application of the pre-trained network is straightforward and user-friendly, which we aspire will enable the community to adopt it as a valuable resource.

The development of large-scale 3D datasets across phylogenetically diverse taxa (e.g., overt [36]) opens up new vistas for comparative research. Likewise, developmental biologists may want to use high-throughput scanning to image hundreds or thousands of specimens as part of an experiment. However, just as DNA sequence data needs bioinformatic algorithms to process massive datasets, large scale image collections require algorithms to digest and extract biologically meaningful data. Algorithms such as this one offer a way forward for powering a “big data” approach to organismal morphology.

## Acknowledgments

This work was supported by funding from the Okinawa Institute of Science and Technology Graduate University (OIST).

The authors would like to gratefully acknowledge H. Skibbe for useful discussion, G. Fischer, F. Hita Garcia, S. Gautam, J. Katzke, F. Azuma and A. Casadei-Ferreira for their help on micro-CT scanning of ants, and A. Buček and T.A. Schoborg for providing fly and termite scans used for the analysis of this project.

